# *Medicago truncatula* possesses a single Shaker outward K^+^ channel: functional characterization and roles *in planta*

**DOI:** 10.1101/720367

**Authors:** Alice Drain, Julien Thouin, Limin Wang, Nicolas Pauly, Manuel Nieves-Cordones, Martin Boeglin, Isabelle Gaillard, Anne-Aliénor Véry, Hervé Sentenac

## Abstract

The model legume *Medicago truncatula* possesses a single outward Shaker K^+^ channel, while *Arabidopsis thaliana* possesses two channels of this type, named SKOR and GORK, the former having been shown to play a major role in K^+^ secretion into the xylem sap in the root vasculature and the latter to mediate the efflux of K^+^ across the guard cell membrane upon stomatal closure. Here we show that the expression pattern of the single *M. truncatula* outward Shaker channel, which has been named MtGORK, includes the root vasculature, guard cells and root hairs. As shown by patch-clamp experiments on root hair protoplasts, besides the Shaker-type slowly-activating outwardly-rectifying K^+^ conductance encoded by *MtGORK*, a second K^+^-permeable conductance, displaying fast activation and weak rectification, can be expressed by *M. truncatula*. A KO mutation resulting in absence of MtGORK activity is shown to weakly reduce K^+^ translocation to shoots, and only in plants engaged in rhizobial symbiosis, but to strongly affect the control of stomatal aperture and transpitational water loss. In legumes, the early electrical signaling pathway triggered by Nod Factor perception is known to comprise a short transient depolarization of the root hair plasma membrane. In absence of MtGORK functional expression, while the rate of the membrane repolarization is shown to be decreased by about 3 times, this defect is without any consequence on infection thread development and nodule production, indicating that the plant capacity to engage rhizobial symbiosis does not require integrity of the early electrical signaling events.

## INTRODUCTION

Potassium (K^+^) can compose up to 10% of the total plant dry weight. This major inorganic constituent of the living cell is the most abundant cation in the cytosol, where it is involved in various functions such as electrical neutralization of negatively charged molecules and control of cell membrane polarization. As an unbound highly mobile abundant osmolyte, K^+^ is also involved in regulation of the cell osmotic potential and related functions such as cell growth or osmotically driven cell and organ movements. It also plays a role in the activation of enzymes, protein synthesis, cell metabolism, and photosynthesis (Clarkson and Hanson, 1980, Nieves-Cordones *et al.*, 2016). Thus, plant growth requires that large amounts of K^+^ ions are taken up by roots from the soil solution and distributed throughout the plant. Several tens of membrane transport systems, which belong to at least 3 families of channels, named Shaker, TPK/KCO and TPC, and 3 families of transporters, named HAK, HKT and CPA, contribute to K^+^ transport (uptake, distribution and compartmentalization) in plants (Mäser *et al.*, 2001; Véry *et al.*, 2014). Among them, the Shaker channel family is the best characterized.

Shaker channels give rise to the main K^+^ conductance of the plasma membrane in most plant cell types (Véry and Sentenac, 2003; Hedrich, 2012). Like their counterparts in animal cells, plant Shaker channels have a tetrameric structure, associating 4 Shaker polypeptides, called alpha-subunits (Daram *et al.*, 1997). A Shaker alpha-subunit consists of a hydrophobic core displaying six transmembrane segments, named S1 to S6, and a pore loop, named P, present between S5 and S6 and carrying the hallmark motif GYGD that plays a central role in the channel selectivity for K^+^. The assembly of the four S5-P-S6 modules in the center of the tetrameric protein structures the K^+^ permeation pathway. Plant Shaker channels, like animal Shakers, are regulated by voltage. The S4 segment harbors positively charged residues (H, R and K) and constitutes the channel voltage sensor. The cytosolic C-terminal part, which begins just after the end of S6, displays a C-linker domain, a cyclic-nucleotide binding domain (CNBD), an ankyrin domain (absent in some alpha-subunits), and a K_HA_ domain rich in hydrophobic and acidic residues (Daram *et al.*, 1997; Nieves-Cordones *et al.*, 2014; see supplementary Figure S1B).

The plant Shaker channel family is strongly conserved, each plant genome harboring about 10 Shaker genes that can systematically be sorted into 5 groups, based on phylogenetic and functional analyses (Véry *et al.*, 2014; see supplementary Figure S1A). Group 1 and 2 members (5 members in Arabidopsis) are characterized as inwardly rectifying channels, mediating K^+^ uptake across the cell membrane. Group 3 channels (a single member of this type in Arabidopsis) display a weak rectification and can thus contribute to both K^+^ uptake and secretion across the cell membrane. Group 4 comprises also a single member in Arabidopsis. It is considered a regulatory subunit since it seems unable to form functional channels by itself but can interact with alpha-subunits from groups 1, 2, and 3 to form heteromeric inward channels with modulated functional features. The last group, group 5, gathers outwardly rectifying channels dedicated to K^+^ secretion from the cell. It comprises 2 members in Arabidopsis, AtSKOR and AtGORK. AtSKOR is strongly expressed in the root vasculature where it plays a major role in K^+^ secretion into the xylem sap and thereby in K^+^ translocation from roots to shoot (Gaymard *et al.*, 1998). *AtGORK* expression has been detected in various tissues and cell types, including root hairs and guard cells. In guard cells, *AtGORK* has been shown to encode the outward conductance that mediate the efflux of K^+^ leading to reduced guard cell turgor upon stomatal closure (Ache *et al.*, 2000; Hosy *et al.*, 2003). In root periphery cells, AtGORK has been shown to mediate an efflux of K^+^ upon the depolarization of the cell membrane that resulted from a strong increase in the external concentration of Na^+^ (Shabala and Cuin, 2007).

So far, besides the work on AtSKOR and AtGORK, very few studies have been aimed at characterizing the functional properties of plant Shaker channels from group 5, and no reverse genetics analysis has highlighted the roles of these functionally-characterized channels (Langer *et al.*, 2002; Sano *et al.*, 2007; Huang *et al.*, 2018). Here we investigate the functional properties and roles of MtGORK, the unique member of the Shaker group 5 in the model legume *Medicago truncatula*.

## RESULTS

### Molecular cloning and primary structure of MtGORK

Phylogenetics analyses indicate that *M. truncatula* Shaker channel group 5 comprises a single member (Damiani *et al.*, 2016a; Wang *et al.*, 2019), Medtr5g077770, hereafter named MtGORK (Supplemental Figure 1A). The corresponding cDNA (2508 pb) was amplified by PCR, allowing sequence analysis of the deduced polypeptide (Supplemental Figures S1B and S1C) and determination of the gene structure (Supplemental Figure S1D). MtGORK possesses the Shaker channel typical hydrophobic core, with the 6 transmembrane segments S1-S6 and the pore loop harboring the GYGD hallmark motif between S5 and S6 (Supplemental Figure S1B and S1C). A C-linker domain, a cyclic nucleotide binding domain, an ankyrin domain and a K_HA_ domain can be identified in the C-terminal region downstream the hydrophobic core, like in the Arabidopsis AtSKOR and AtGORK outward channels. The percentages of identity and similarity between MtGORK and AtSKOR or AtGORK are close to 70% and 85%, respectively. The two residues of the P loop and 2 residues of the S6 transmembrane segment that have been identified in AtSKOR as contributing to the dependency of the channel voltage-sensitive gating on the external concentration of K^+^ (Johansson *et al.*, 2006), conserved in AtGORK, are also present in MtGORK (Supplemental Figure S1C).

### Functional characterization in *Xenopus* oocytes

Depolarization of the membrane elicited an outward current in oocytes injected with *MtGORK* cRNA and not in control oocytes injected with water (Figure 1A). The exogenous macroscopic current displayed slow sigmoidal activation kinetics and reached a steady-state value within *ca.* 2 s (Figure 1A). Steady-state I-V curves displayed a strong outward rectification (Figure 1B). Comparison of the I-V curves obtained in presence of 10, 30 or 100 mM K^+^ in the external solution revealed that increasing this concentration resulted in a positive shift of the activation potential threshold, *i.e.*, the threshold beyond which, when the potential was shifted to more positive values, outward currents became detectable (Figure 1B). The experimental curves describing the dependency of the channel relative open probability, Po/Po_max_, on voltage (obtained from MtGORK deactivation currents recorded at + 50 mV after pre-pulses varying from −100 mV to +80 mV) in presence of 10, 30 or 100 mM external K^+^ were fitted with the classical two-state Boltzmann law (Ache *et al*., 2000). The results indicated that the channel half activation potential (*Ea50*: membrane potential at which the channel relative open probability is 0.5) was strongly dependent on external K^+^ concentration, being shifted by +50 mV when this concentration was increased from 10 to 100 mM (Figure 1C). Such a regulation by external K^+^ ensures that the outward rectification of MtGORK is total regardless of the concentration of K^+^ prevailing outside, and thus that this channel is strictly dedicated to K^+^ release. Besides the sensitivity to voltage and external K^+^, MtGORK was also found to be sensitive to the external pH, the outward current being decreased by about 50% when the pH was decreased from 7.5 to 5.6 (Figure 1D). Thus, the pH sensitivity of MtGORK appears stronger than that reported in AtGORK (Ache *et al.*, 2000).

**Figure 1.**
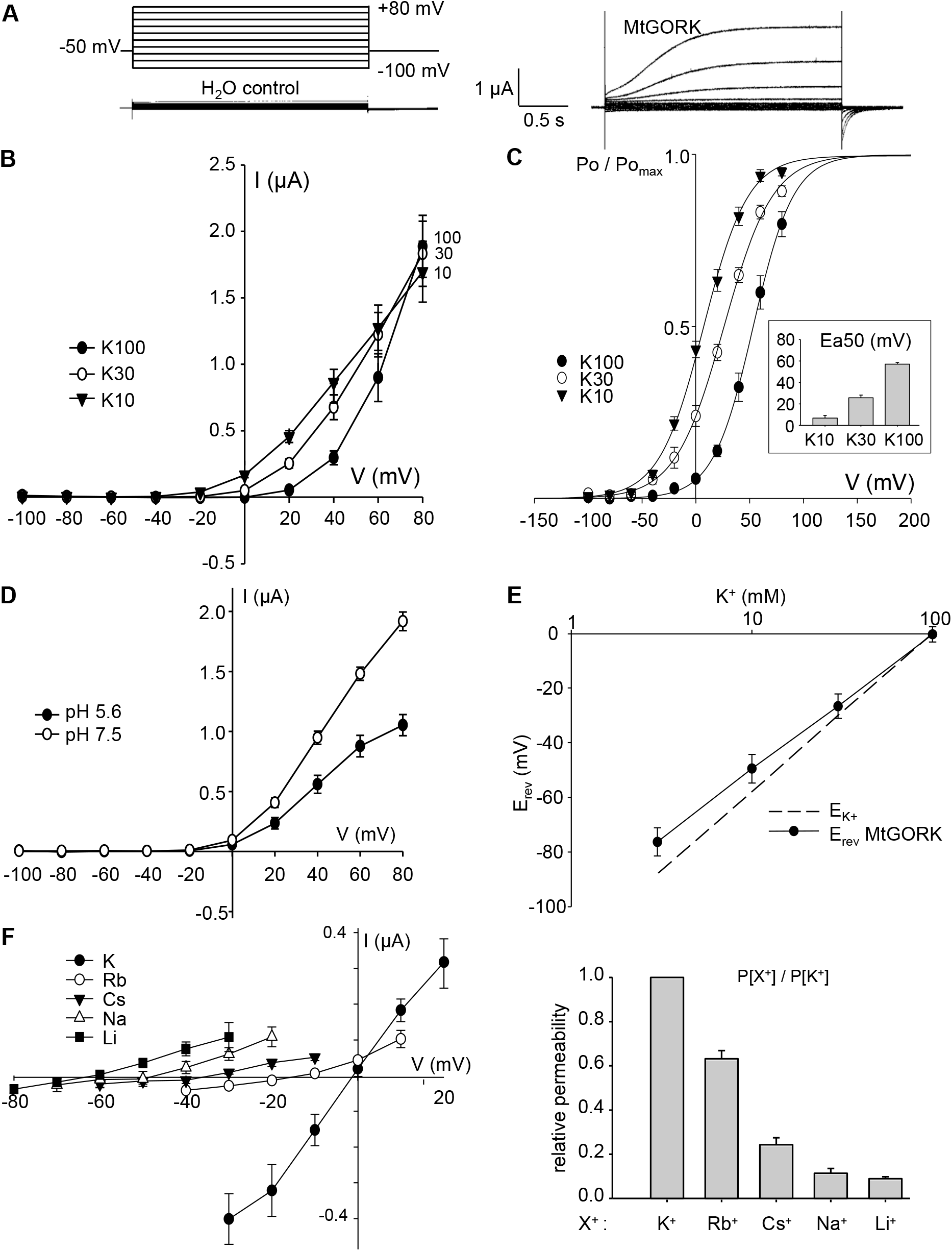
Functional characterization of MtGORK in *Xenopus* oocytes. (A) Voltage clamp protocol (top left) and typical currents recorded in control oocytes injected with H_2_O (bottom left) or injected with *MtGORK* cRNA (right) in 100 mM K^+^ solution. Voltage-clamp pulses varied from −100 to +80 mV, in increments of 20 mV. Every episode of imposed voltage lasted 3 seconds. The holding potential was −50 mV. (B) MtGORK current-voltage relationships at different external K^+^ concentrations: 10, 30 or 100 mM. Mean ± SE, n ≥ 8. (C) Effect of the membrane voltage and the external K^+^ concentration on MtGORK open probability. The relative open probability (Po/Pomax) was obtained from the analysis of deactivation currents upon return to the holding voltage after the activation pulse (mean ± SE, n ≥ 6). The solid line represents Boltzmann fits to the mean Po/Pomax values. The mean values (± standard error, n ≥ 6) of the half-activation potential (Ea50) of the MtGORK channel obtained from these fits in the different concentrations of K^+^ are presented in the inset. (D) Activation of MtGORK currents by external alkalinization. The external solution contained 100 mM K^+^. The external pH was 5.6 or 7.5. Means ± SE, n = 4. (E) Variation of MtGORK reversal potential of currents (Erev) with the external concentration of K^+^ (mean ± SE, n ≥ 8). Erev was determined in each solution using a tail-current protocol: After activation of MtGORK channels at + 60 mV, voltage pulses were performed at lower voltages flanking Erev, and Erev was obtained from the analysis of the deactivation currents. The dashed line indicates K^+^ equilibrium potential. (F) Permeability of MtSKOR to different monovalent cations. (Left) MtGORK deactivation currents were recorded using a tail-current protocol (as above) in bath solutions containing 100 mmole.l^−1^ of either K^+^, Rb^+^, Cs^+^ Na^+^ or Li^+^. Mean deactivation currents ± SE, n≥ 6. (Right) Permeability ratios of the different cations with respect to that of K^+^, calculated (from the variations of Erev) using the Goldman-Hodgkin-Katz equation. Mean ± SE, n≥ 6.

A last series of experiments was aimed at characterizing the ionic selectivity of MtGORK. As expected, the current reversal potential, *E*_*rev*_, determined from classical analysis of tail current recordings (Ache *et al.*, 2000), was found to be dependent on the external concentration of K^+^ (Figure 1E). *Erev* shifted by about +50 mV for a 10-fold increase in external K^+^ concentration, indicating that MtGORK displays a strong selectivity for K^+^ (a channel exclusively permeable to K^+^ would have given rise to a shift of *ca*. +58 mV). Shifts in *E*_*rev*_ were also recorded upon replacement of K^+^ in the external medium by another alkali cation, either Li^+^, Na^+^, Rb^+^ or Cs^+^, at the same concentration (100 mM) (Figure 1F). In such experiments, the magnitude of the *E*_*rev*_ shift reflects the relative permeability of the substituting cation and allows to determine this permeability, reported to that to K^+^, using the so-called Goldman equation (Hille, 2001). MtGORK displayed the following permeability sequence (Eisenman’s series IV; Eisenman, 1961), K^+^>Rb^+^>Cs^+^>Na^+^≈Li^+^ (Figure 1F), which is identical to that reported for AtSKOR (Gaymard *et al.*, 1998). Like in AtSKOR, the relative permeability to Rb^+^ is rather high (*ca.* 0.6), while that to Na^+^ is weak (<0.1).

### A KO mutation in *MtGORK* results in absence of Shaker type outward K^+^ conductance in guard cells

A *M. truncatula* line (cv Jemalong A17) named NF9352, displaying an insertion of the *Tnt-1* retrotransposon in the first exon of *MtGORK* (Supplemental Figure S1D) was obtained from the mutant collection of the Noble Foundation and self-pollinated to produce the F2 generation. The plant that was amplified possessed this mutation, hereafter named *mtgork*, at the hemizygous state. Genotyping experiments (PCR) were carried out to identify both mutant plants homozygous for the *mtgork* mutation and control WT plants possessing a wild type (WT) genotype at this locus. RT-PCR experiments could not amplify *bona fide MtGORK* transcripts in mutant plants homozygous for the *mtgork* mutation (Figure 2A), providing evidence that *mtgork* is a knock-out (KO) mutation. The selected mutant and WT lines were further amplified for phenotyping experiments. Visual observations of the plants homozygous mutant or WT for the *mtgork* mutation did not allow to detect any specific phenotype. Furthermore, measurements of root and shoot biomass production in plants grown in different conditions (*in vitro*, in greenhouse or growth chamber on compost or sand-vermiculite mixture) at different growth stages and inoculated with a rhizobial strain (*S. meliloti* 1021 strain) or not inoculated did not reveal any specific defect in plant development (Supplemental Figure S2).

**Figure 2.**
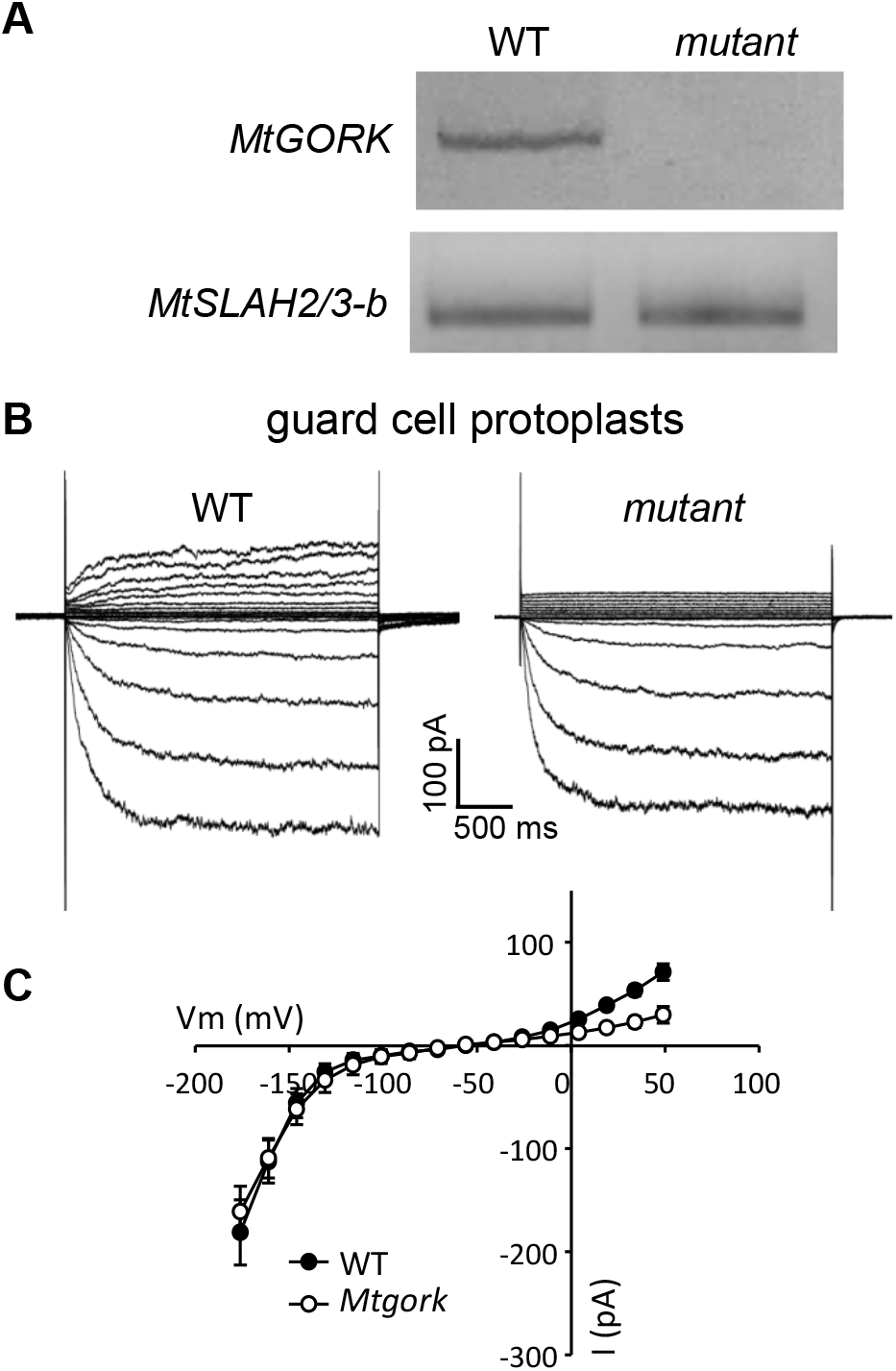
The *mtgork* mutation results in absence of MtGORK functional expression in guard cells. (A) PCR analyses reveal that mutant plants homozygous for the *mtgork* mutation do not express *MtSKOR* transcripts. Plants homozygous for the *mtgork* mutation (mutant) or displaying a wild type genotype for this mutation (WT) were grown *in vitro* on Fahraeus medium for 10 days before RNA extraction from whole plants. PCR experiments did not detect *MtGORK* transcripts in the mutant plants. The gene *MtSLAH2/3-b* (*Medtr6g045200*) (Damiani *et al.*, 2016a) was taken as control (lower panel). (B and C) Comparison of K^+^ currents in guard cell protoplast from WT and mutant plants. Currents were recorded using the patch-clamp technique in the whole-cell configuration. The ionic composition of the bath solution was 10 mM K-glutamate, 10 mM CaCl_2_ and 10 mM Mes-Tris, pH 5.8. Patch-clamp pipette solution: 100 mM K-glutamate, 5 mM EGTA, 1 mM CaCl_2_ (free Ca^2+^ = 20 nM), 2 mM MgCl_2_, 2 mM Mg-ATP, 10 mM Hepes-Tris, pH 7.5. The osmolarity of bath and pipette solutions was adjusted to 480 and 500 mosmol/Kg with D-mannitol. “Outward” whole-cell currents were recorded applying successive pulses of clamped voltage from −71 to 49 mV (after liquid junction potential (LJP) correction) in 15 mV increment from a holding potential at −71 mV; inward currents were recorded applying voltage pulses from −71 to −161 mV in −15 mV increment from a holding potential at −71 mV. (B): Representative current traces recorded in WT and mutant protoplasts. (C): Steady state current-voltage (I-V) plot. Means ±SE (WT: n = 5; mutant: n = 9).

Patch-clamp experiments carried out on guard cell protoplasts revealed that homozygous plants for the *mtgork* mutation did not display the typical Shaker-like outward K^+^ currents that were recorded in WT plants (Figure 2B). Small instantaneously-activating outward K^+^ currents were detected in the mutant protoplasts (Figure 2B & C). On the other hand, the inward K^+^ currents were very similar in mutant and WT protoplasts. Based on their activation kinetics and current-voltage curve, they are likely to be mediated by inwardly rectifying Shaker channels like in Arabidopsis guard cells (Lebaudy *et al.*, 2008).

### The *mtgork* KO mutation results in impaired control of transpirational water loss

A sharp reduction in the rate of leaf transpirational water loss rapidly (<20 min) occurred in WT plants after leaf excision, whereas no significant change in this rate could be observed in the mutant plants over 60 min. Thus, as a predicted consequence of the absence of Shaker-like outward conductance in guard cells, a defect in the capacity of stomatal closure was displayed by the mutant plants (Figure 3). The dotted lines plotted in Figure 3 correspond to data obtained in a similar experiment in Arabidopsis, with mutant plants that did not express the *AtGORK* gene shown to encode the guard cell outward Shaker conductance (Hosy *et al.*, 2003).

**Figure 3.**
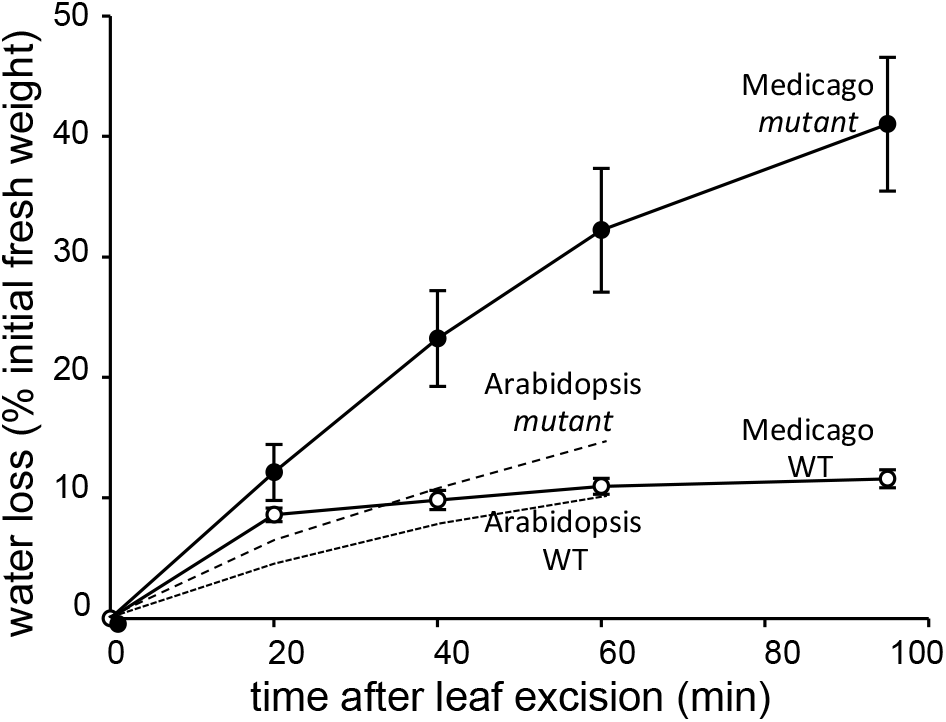
The *mtgork* mutation results in impaired stomatal closure. Plants homozygous for the *mtgork* mutation or displaying a wild type genotype for this mutation were grown on compost in greenhouse for 6 weeks. Leaves were excised during the light period and water loss was determined by monitoring the decrease in fresh weight of the excised leaves. Means ± SE, n = 3.

### *MtGORK* is expressed in root stellar tissues and root hairs

*MtGORK* expression was first investigated *in silico* using public databases [“eFP Browser Medicago”, (http://bar.utoronto.ca/), and “Medicago Gene Atlas”, (http://mtgea.noble.org/v3/)]. The data indicated that *MtGORK* is expressed in leaves, stem, roots and nodules. Then, analysis of *M. truncatula* roots transformed with a *MtGORK promoter-GUS* reporter gene construct indicated that the expression pattern of *MtGORK* includes root hairs, root vascular tissues and nodule vasculature (Figure 4). Evidence for the expression of *MtGORK* in root hairs has also been provided by RNA-Seq analyses (Damiani *et al.*, 2016a).

**Figure 4.**
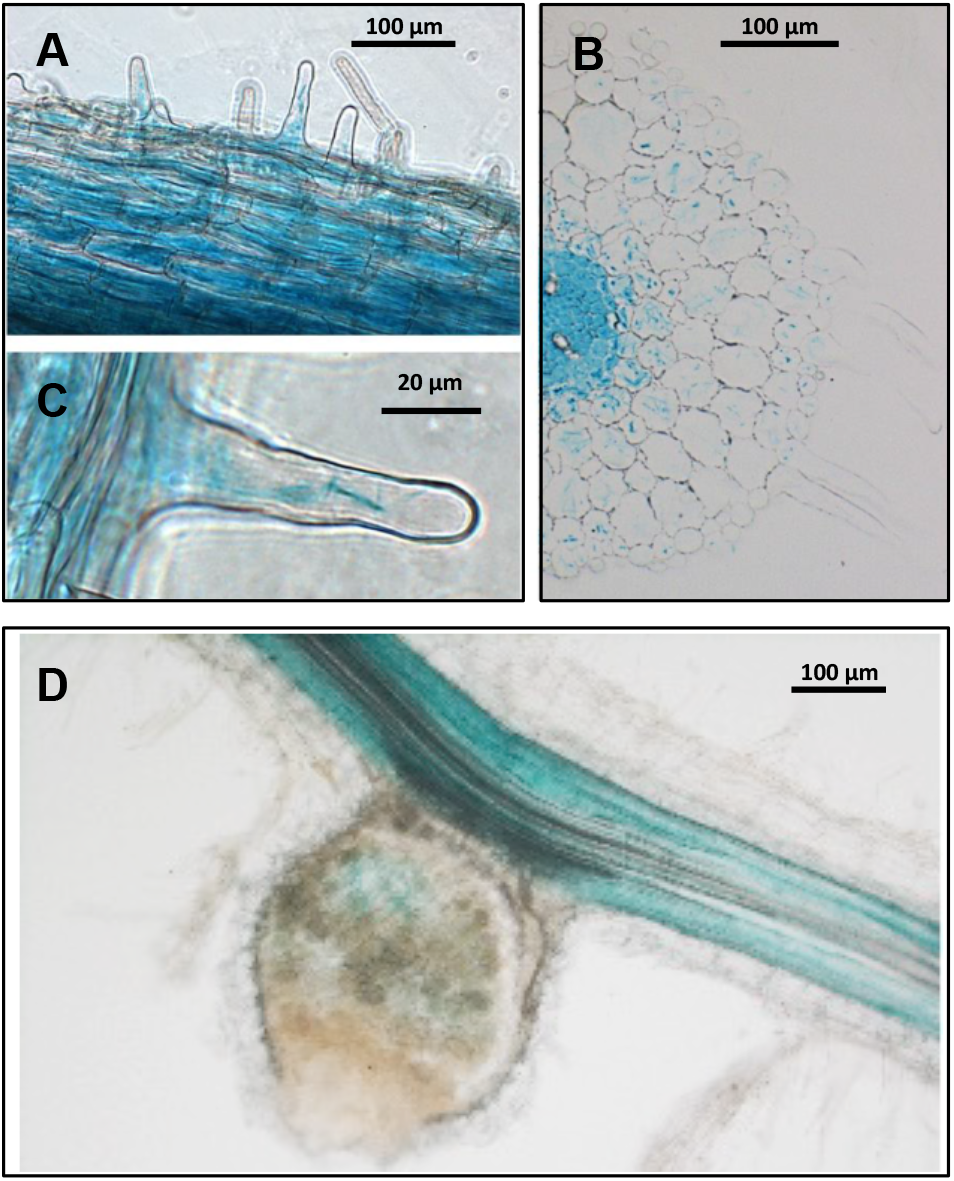
Expression pattern of the *MtGORK* gene in roots. (A) Representation of the *promoter-GUS* reporter gene transformed in *M. truncatula* roots using *A. rhizogenes*. (B) Histochemical analysis of *MtGork* expression in *Medicago truncatula* roots GUS activity in *M. truncatula* composite plants expressing *pMtGork*::GUS fusion. *MtGork* is expressed in root epidermis (a), in root hairs (b), in root stellar tissues (c) and in root nodules (d).

In Arabidopsis, the outward Shaker *AtGORK* is expressed in root hairs and *AtSKOR* in the root stele. We have investigated the consequences of the *mtgork* mutation in these root tissues/cell types through the analysis of (i) the translocation of K^+^ towards the shoots, (ii) the electrical properties of the root hair cell membrane, and (iii) the early electrical signal induced by Nod factor (NF) perception in root hairs and the plant capacity to engage rhizobial symbiosis.

### K^+^ translocation towards the shoots

Experiments carried out to assess the contribution of MtGORK to K^+^ translocation to shoots revealed that the *mtgork* mutation poorly affected the shoot K^+^ content. A reduction in this content could be observed in inoculated plants, but it was slight (*ca.* 13%), and no statistically significant difference was observed in non-inoculated plants (Figure 5). In agreement with these results, the absence of MtGORK activity was found to be without any significant effect on the concentration of K^+^ in exuded xylem sap and on the volume of exuded sap (due to root pressure) upon shoot excision, whatever the plant status, inoculated or non-inoculated (Figure 6).

**Figure 5.**
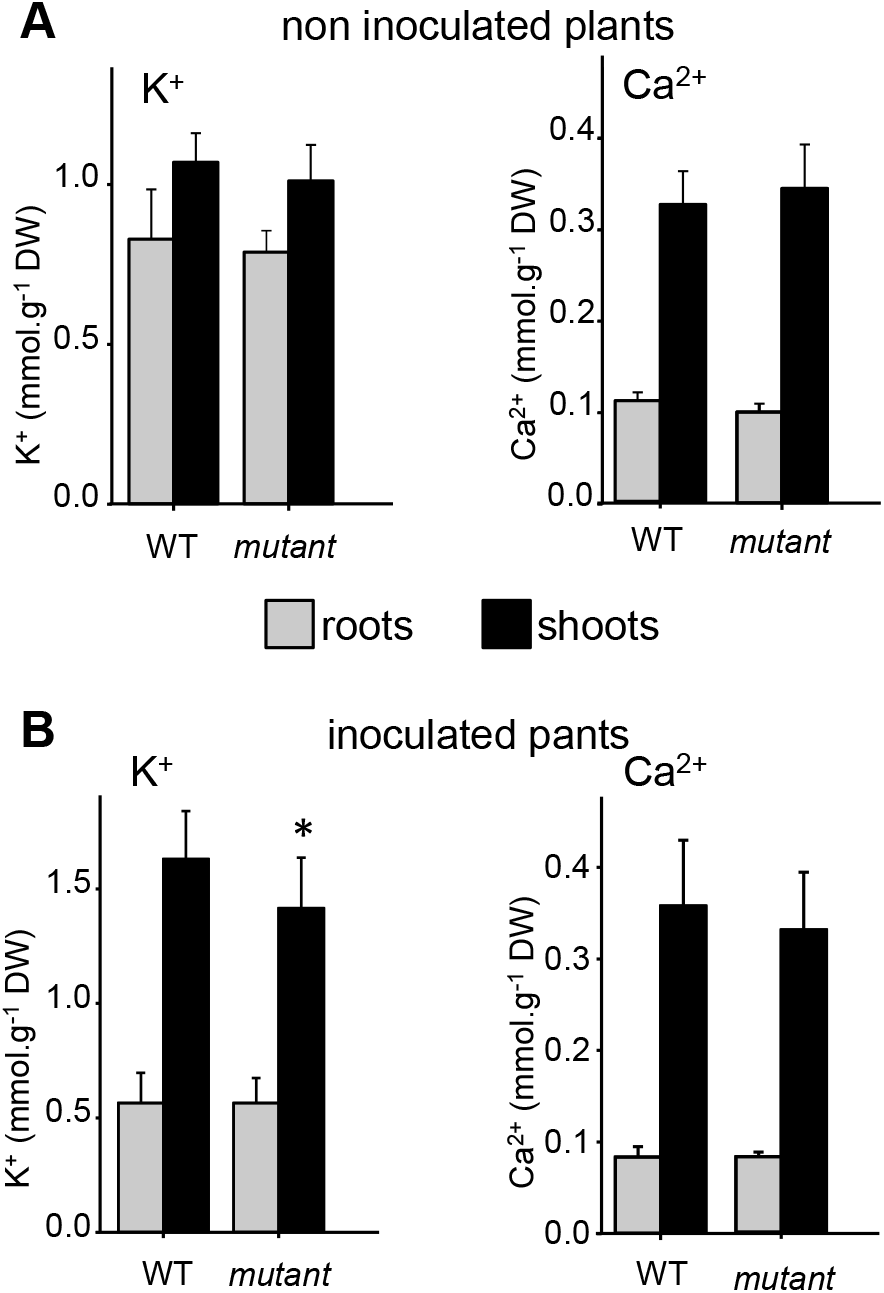
Effect of the absence of MtGORK activity on root and shoot K^+^ contents in non-inoculated and inoculated plants. Plants homozygous for the *mtgork* mutation (mutant) or displaying a wild type genotype for this mutation (WT) were transferred onto sand-vermiculite mixture 3 days after germination. Shoots and roots were collected for K^+^ and Ca^2+^ assays after 8 weeks of growth. Means ± SE; n = 12. (A) Non inoculated plants. (B) Inoculated plants. Inoculation was performed with *S. meliloti* strain 1021 at the end of the second week of growth.

**Figure 6.**
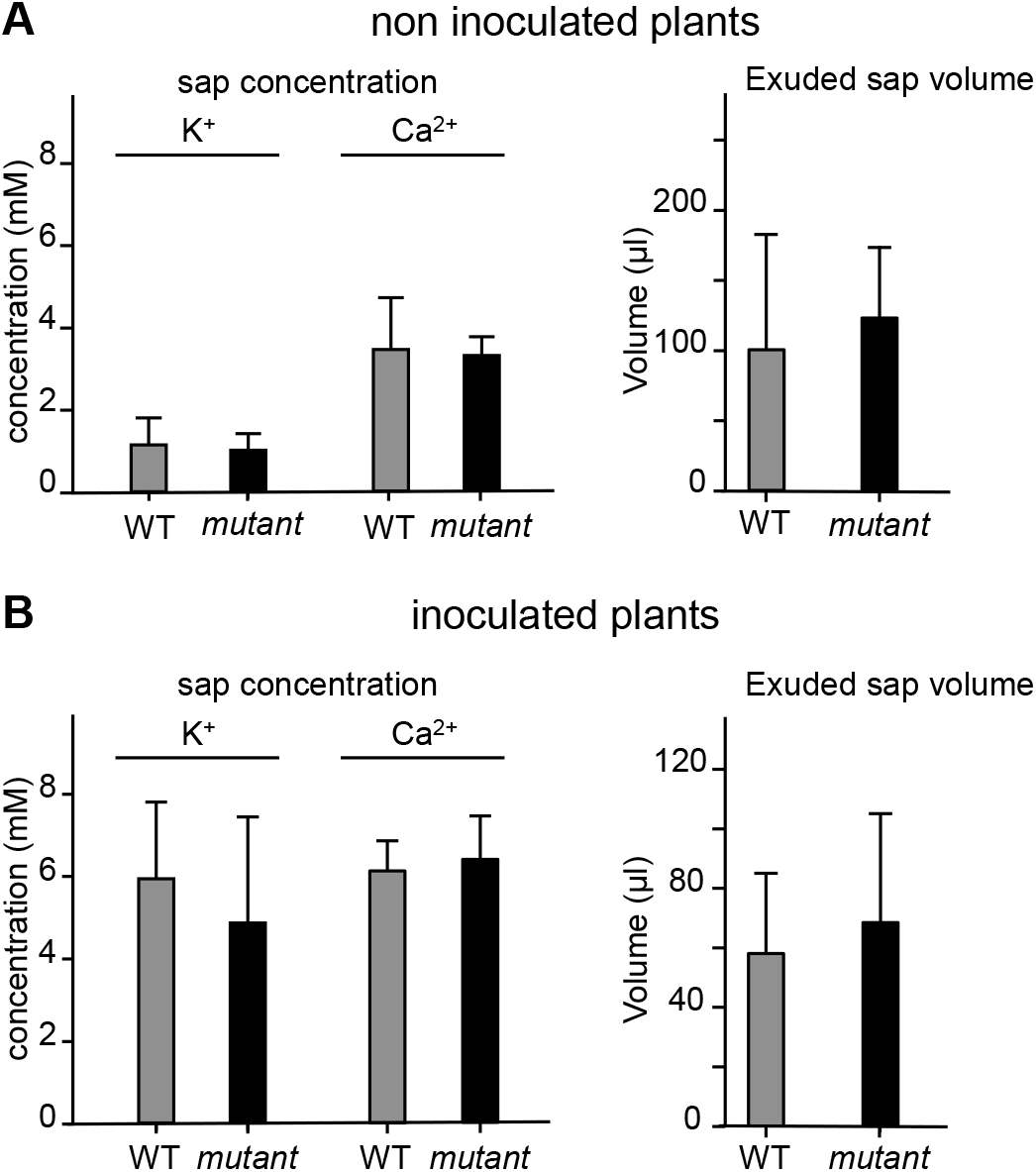
Absence of MtGORK activity does not impact the xylem flux of K^+^ from roots to shoot. Plants homozygous for the *mtgork* mutation (mutant) or displaying a wild type genotype for this mutation (WT) were grown as described in the legend to Figure 5. After 8 weeks of growth, shoots were excised at about 1 cm above the soil. About 30 min after shoot excision, exsuded sap was collected for 24 h. The volume and K^+^ and Ca^2+^ concentrations of the exuded sap were measured. Means ± SE; n = 12. (A) Non inoculated plants. (B) Inoculated plants. Inoculation was performed with *S. meliloti* strain 1021 at the end of the second week of growth.

### Electrical properties of the root hair plasma membrane

Patch-clamp recordings in root hair protoplasts from WT plants revealed that two distinct types of outward K^+^ conductances (Figure 7A; upper panels), differing at least in their activation kinetics and in the shape and size of the deactivation current, as previously reported (Wang *et al.*, 2019), could dominate the membrane permeability to K^+^. Here, in experiments made in a native context, we use the term "conductance" to refer to a type of permeation pathway that can be mediated by a single molecular identity or by a set of channels of several identities but displaying similar properties.

**Figure 7.**
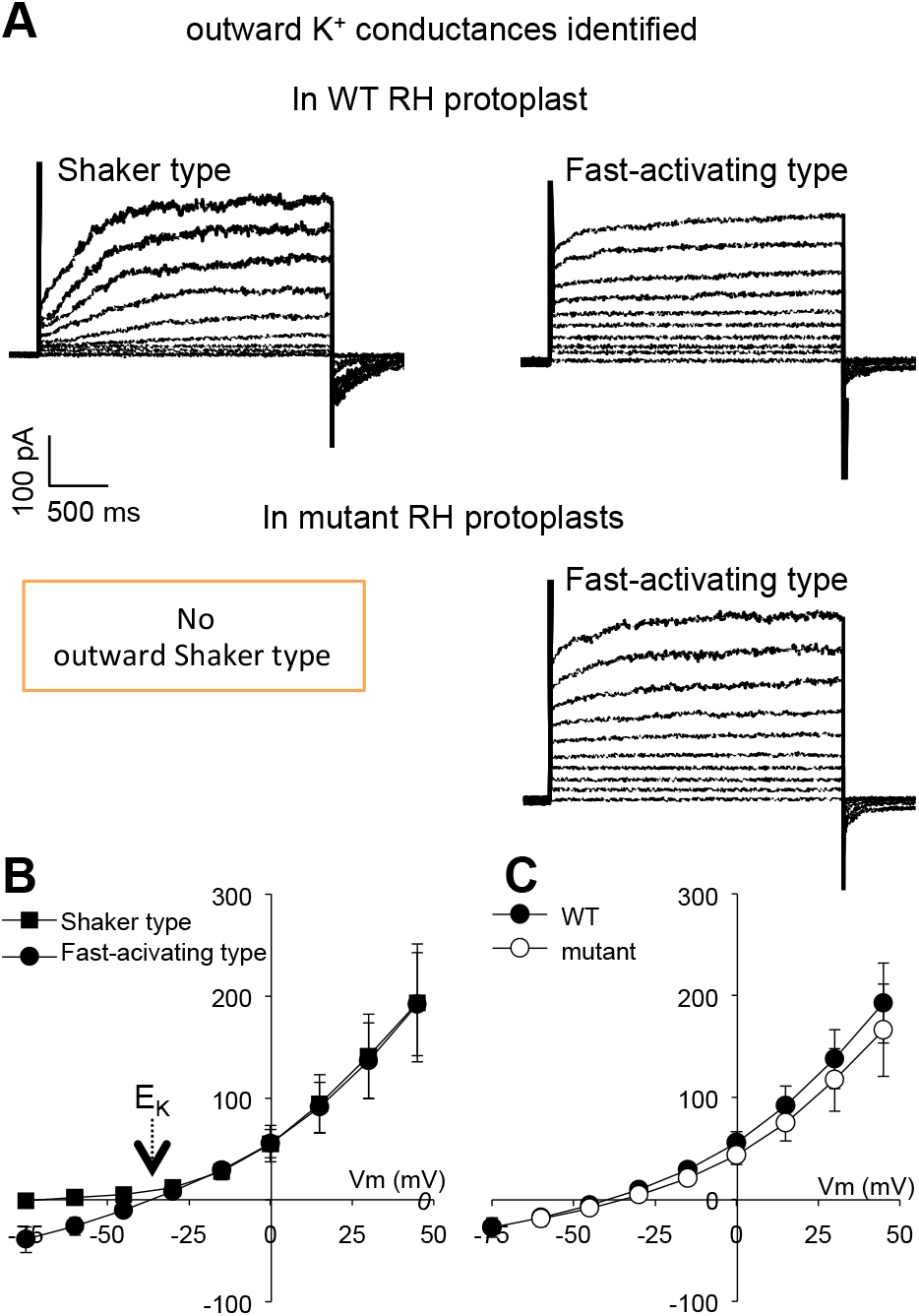
Outward K^+^ currents in root hair protoplasts from WT or mutant plants. Root hair (RH) protoplasts were enzymatically obtained from plants homozygous for the *mtgork* mutation (mutant) or displaying a wild type genotype for this mutation (WT). The bath solution contained 30 mM K-gluconate, 1 mM CaCl_2_, 10 mM Mes-Tris, pH 5.6. Patch-clamp pipette solution: 150 mM mM K-gluconate, 10 mM EGTA, x mM CaCl_2_ (à voir par Anne), 2 mM MgCl_2_, 2 mM Mg-ATP, 10 mM Hepe-Tris, pH 7.4. The osmolarity of the bath and pipette solutions was adjusted to 290 and 300 mosmol/Kg respectively, with D-sorbitol. Voltage clamp proyocol: pulses from −75 to 60 mV (after correction of LJP), 15 mV increment, and holding potential at −75 mV. (A) Representative current traces recorded in WT and mutant protoplasts. in WT protoplasts (top panels), two types of macroscopic conductances could be identified: the top left recording is typical of outward Shaker channels (slow and sigmoidal activation), and the top right example can be ascribed to the fast-activating outward cationic conductance previously described (Wang *et al.*, 2019). In mutant protoplasts, the outward Shaker conductance was not observed, while the fast-activating cationic outward conductance was present. (B) Steady state current-voltage (I-V) relationship of the outward Shaker type conductance and of the fast-activating outward cationic conductance derived from patch-clamp recordings in 28 WT protoplasts. The membrane outward conductance to K^+^ was dominated by the outward Shaker type conductance in 10 protoplasts, and by the fast-activating cationic conductance in 13 protoplasts. In 5 protoplasts, a contribution of both conductances to the outward current could be distinguished. I-V curves were derived for the 10 protoplasts dominated by the Shaker-type conductance and the 13 protoplasts dominated by the fast-activating conductance. Means ± SE. (C) Steady state current-voltage relationship of the membrane outward conductance in WT and mutant protoplasts. Means ± SE. WT: n = 28 (same protoplasts as in panel B). Mutant: n = 15.

In some protoplasts, the dominant conductance displayed a slow sigmoidal activation of currents and slow current deactivation kinetics upon return to the holding voltage (Figure 7A, left panel), reminiscent of the features of MtGORK when expressed in *Xenopus* oocytes (Figure 1C) or when characterized *in situ* at the guard cell membrane in WT plants (Figure 2B, left panel), and typical of outward Shaker channels (Gaymard *et al.*, 1998; Ache *et al.*, 2000; Langer *et al.*, 2002; Sano *et al.*, 2007; Huang *et al.*, 2018). Such a conductance has also been recorded in root hairs from Arabidopsis (Ivashikina *et al.*, 2001) and *Medicago sativa* (Bouteau *et al.*, 1999). On the other hand, this type of conductance was not observed in mutant plants homozygous for the *mtgork* mutation (Figure 7A, lower left panel).

The second conductance that could be the dominant one in WT protoplasts (Figure 7A, upper right panel) was also found in *mtgork* mutant root hair protoplasts (Figure 7A, lower right panel). It is hereafter named fast-activating outward cationic conductance as in Wang *et al.* (2019) since it displays a more rapid activation than that of MtGORK. The recorded current traces reveal an instantaneous component and a time-dependent component, the relative part of the latter increasing with depolarization (Figure 7A). The deactivation currents observed upon return to the holding voltage were small when compared with those of MtGORK. The fact that the time-dependent component of this second conductance was also displayed by the current traces obtained in mutant protoplasts that did not express MtGORK, indicates that this conductance does not result from the addition of a MtGORK component to an instantaneously-activating conductance.

In a few WT protoplasts (5 out of 28), the recorded traces indicated that the above described conductances were simultaneously active at the root hair cell membrane: *e.g*., a large instantaneously activating component could be distinguished together with large and slowly deactivating tail currents. In most cases however, the outward conductance of the membrane to K^+^ could be considered as essentially resulting from the activity of a single conductance type, either the fast-activating conductance or the MtGORK one, these two situations having rather similar frequencies: the former conductance was the dominating one in 13 protoplasts, while the MtGORK conductance dominated in 10 protoplasts. Sorting the recorded protoplasts into these two categories and deriving the corresponding current-voltage curves revealed that the protoplasts dominated by the MtGORK conductance displayed a strong outward rectification, reminiscent of that obtained in oocytes expressing MtGORK, in contrast to the protoplasts dominated by the fast-activating conductance (Figure 7B). Finally, the current-voltage curve derived for this set of 28 WT protoplasts, treated as a whole, was similar to the one obtained for 15 protoplasts from mutant plants homozygous for the *mtgork* mutation in terms of current magnitude (Figure 7C). Altogether, these results suggest that the absence of MtGORK conductance in root hairs of the mutant plant was compensated by an increase in the expression or activity of the fast-activating outward conductance.

### MtGORK contributes to repolarization of the root hair cell membrane following Nod Factor induced depolarization

Initiation of symbiotic interactions with N_2_-fixing rhizobia in legumes is triggered at the root hair cell membrane in response to nodulation factors (NF) secreted by rhizobia, and involves complex signaling events (Felle *et al.*, 1998; Oldroyd and Downie, 2008). The earliest events that have been reported, together with ROS production (Puppo *et al.*, 2013; Damiani *et al.*, 2016b), are changes in ion fluxes, H^+^, Ca^2+^, anion (Cl^−^) and K^+^, through the root hair plasma membrane resulting in a transient depolarization of this membrane (Felle *et al.*, 1998). Continuous recordings of the local concentrations of Ca^2+^, H^+^, Cl^−^ and K^+^ at the root surface using extra-cellular ion selective micro-electrodes and of the changes in membrane potential using an intracellular micro-electrode have shown that NF perception rapidly results (within *ca.* 1 min) in an increase in net Ca^2+^ influx, followed by a net efflux of anions and possibly an inhibition H^+^-excretion. Altogether, these events result in a strong membrane depolarization, which activates voltage-sensitive K^+^ channels, allowing an efflux of K^+^ that repolarizes the membrane, a process to which an activation (or a re-activation) of H^+^ excretion by proton pumps could contribute (Felle *et al.*, 1998). The hypothesis that MtGORK contributes to the efflux of K^+^ involved in the repolarization of the membrane during this action potential-like signaling process was tested by comparing the kinetics of membrane repolarization, recorded by microelectrode impalement, in WT and mutant plants as described in Figure 8. Thereby, the rate of repolarization (mV.min^−1^) was found to be about two times slower in mutant plants homozygous for the *mtgork* disruption when compared with the control WT plants (Figure 8C).

**Figure 8.**
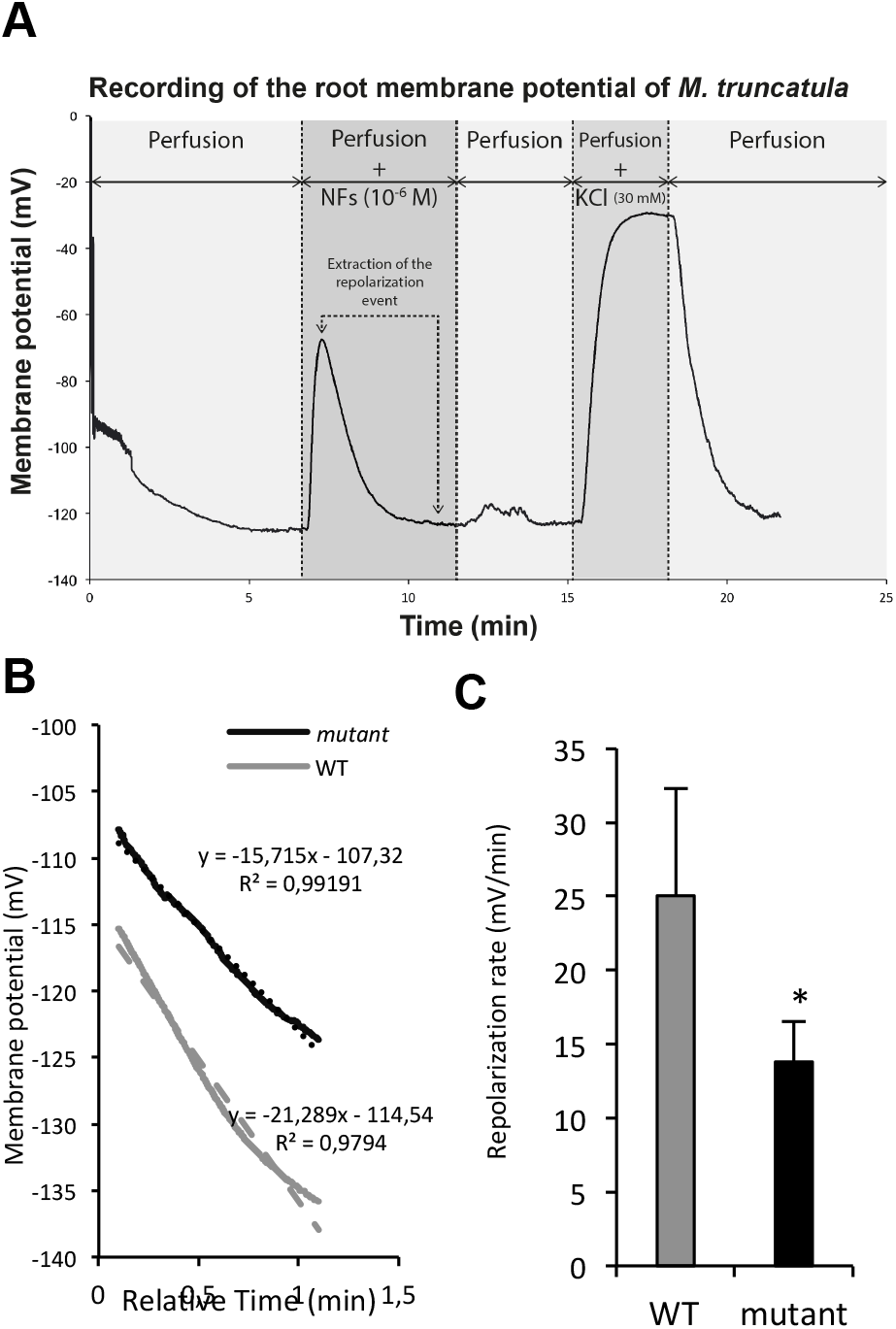
Role of MtGORK in repolarization of the root hair cell membrane after the initial depolarization induced by Nod Factor perception. (A) Recording of the membrane potential variations induced by NF factor treatment in a plant displaying a wild type genotype (WT) for the *mtgork* mutation. The external solution bathing the root (bath background solution: bbs) when the micropipette was impaled in an epidermal cell of the root hair zone contained 0.1 mM KCl, 0.1 mM CaCl_2_, 0.1 mM NaCL and 2 mM MES-Bis-Tris-Propane, pH 6.5. When a stable signal was observed, the bbs was replaced by Nod Factor (NF) solution, which contained 10^−6^ M Nod Factor in bbs. The treatment resulted in a rapid depolarization of the cell membrane followed by a repolarization to the initial value while NFs were still present in the percolated solution. Then, the NF solution was replaced by bbs, allowing to check that the membrane potential remained close to the initial value recorded before the NF treatment. The bbs was thereafter replaced by 30 mM KCl (in bbs) to check the depolarizing effect of a high K^+^ concentration. The 30 mM KCl solution was thereafter replaced by bbs to check whether the impaled cell could restore its membrane potential to the initial value. When this test was positive, the recording was used for analysis of the repolarization phase during the NF treatment. (B) Analysis of the repolarization phase. The apparently linear part of the repolarization phase (lasting about 1 min, from *ca.* 0.1 min after the beginning of this phase) was extracted and fitted with a linear regression to derive the mean slope in mV.min^−1^) of the recording, taken as an estimate of the repolarization rate. Black and grey curves: example of recordings obtained in a plant homozygous for the *mtgork* mutation (mutant) or displaying a wild type genotype for this mutation (WT), respectively. (C) Repolarization rates in WT and mutant plants. Means ± SD, n = 5. The asterisk indicates that the difference is statistically significant (Student test, P<0.05).

### MtGORK activity is not necessary for infection thread development and nodule formation

The number of infection threads developed in plants grown *in vitro* on Farhäeus medium and observed at either 3 or 5 days after root inoculation (dpi) with *S. meliloti* was not significantly different between mutant plants homozygous for the *mtgork* disruption and control WT plants (Figure 9A). In agreement with this result, when growth continuously occurred *in vitro* on Farhäeus medium, the number of nodules determined at either 7, 14 or 21 dpi was similar in the mutant and WT plants (Figure 9B).

**Figure 9.**
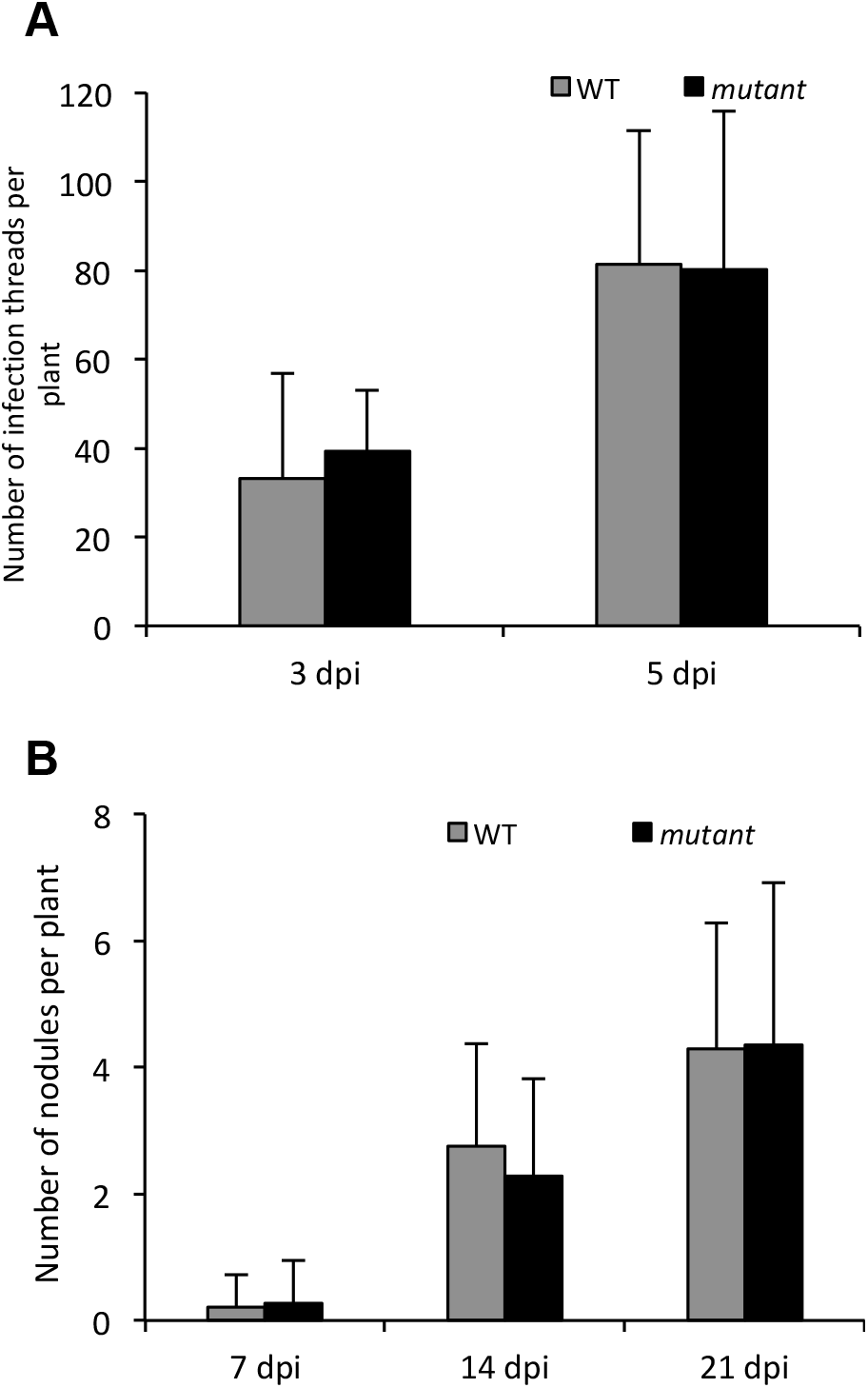
*In vitro* capacity of the plant to produce infection threads in presence of rhizobia and to develop nodules is not impaired by absence of MtGORK activity. Germinated seedlings homozygous for the *mtgork* mutation (mutant) or displaying a wild type genotype for this mutation (WT) were transferred onto Fahraeus agar medium in Petri dishes and inoculated with *S. meliloti* two days later. (A) Number of infection threads observed at 3 and 5 days post-infection (dpi). The inoculated *S. meliloti* strain was RM2011 lacZ, and the collected roots were stained to reveal LacZ activity for easier detection of infection threads. Means ± SE, n = 10. Statistical analysis (Tukey test at the 5% level) indicates that there was no significant difference between the two types of plants. (B) Number of nodules at 7, 14 and 21 dpi. The inoculated *S. meliloti* strain was 1021 DsRED, and the collected roots were observed using DsRED fluorescence microscopy, allowing easier detection of nodules. Means ± SE, n = 24. No statistically significant difference between the two types of plants (Tukey test at the 5% level).

## DISCUSSION

### A single outwardly rectifying Shaker channel in *M. truncatula*

The Shaker channel family comprises a single outwardly rectifying member in *M. truncatula* (Suplemental Figure S1A). It is interesting to note that *in silico* analysis of the genome sequence of *Lotus japonicus* (https://lotus.au.dk/) reveals a single outwardly rectifying member in the Shaker family of this legume model too. In Arabidopsis, the Shaker gene family comprises 2 genes coding for outwardly rectifying channels, *AtGORK* and *AtSKOR*, and 7 genes for inwardly (or weakly-inwardly) rectifying channels (Supplemental Figure S1A). The phylogenetic structure of the Shaker family is strongly conserved in plants: about 10 genes and always 5 subgroups, the genes coding for inwardly-rectifying channels (groups 1 to 4) being more numerous than those coding for outwardly-rectifying channels (group 5) (Véry *et al.*, 2014).

Analyses of the functional properties of plant inwardly rectifying channels in heterologous systems have revealed distinctive features and regulations, in terms of voltage sensitivity, affinity for external K^+^ or sensitivity to pH (Véry and Sentenac, 2003; Véry *et al.*, 2014). Formation of heteromeric channels associating subunits encoded by different inward Shaker genes can further increase this functional diversity (Reintanz *et al.*, 2002; Duby *et al.*, 2009; Jeanguenin *et al.*, 2011). The corresponding information presently available on outwardly rectifying channels is much more reduced. Besides MtGORK and the Arabidopsis AtGORK (Ache *et al.*, 2000; Hosy *et al.*, 2003) and AtSKOR (Gaymard *et al.*, 1998; Johansson *et al.*, 2006), only 3 other outward K^+^ channels have been characterized at the functional levels in heterologous systems, NTORK from tobacco (Sano *et al.*, 2007), PTORK from poplar (Langer *et al.*, 2002) and CmSKOR from melon (Huang *et al.*, 2018). These outward Shaker channels share a common functional feature: their activation threshold depends on the external concentration of K^+^. This regulation by external K^+^, also observed in outward K^+^ conductances recorded *in planta* (Schroeder, 1989; Blatt, 1991; Blatt and Gradman, 1997; Roelfsema and Prins, 1997; Wang *et al.*, 2019), ensures that the channels open only when the transmembrane K^+^ electrochemical gradient is outwardly directed, regardless of the external concentration of K^+^, and thus that these channels are dedicated to the function of K^+^ secretion into the apoplasm/external medium. Dominated by this tight regulation, the functional diversity of outward Shaker channels in plants seems to be rather reduced, when compared to that of inward Shaker channels. This might explain that the number of outward Shaker genes is low in every plant genome.

### Both MtGORK and a non-Shaker K^+^-permeable outward conductance can be expressed in root hairs

Differences in the current activation kinetics and in the shape and size of deactivation currents displayed by protoplasts from wild-type plants supported the hypothesis that two types of K^+^-permeable outward conductance could be active at their plasma membrane. These two types of conductance do not correspond to two different cell types, *e.g.* trichoblasts *versus* atrichoblasts or cortical cells, since they have also been observed in spheroplasts obtained from the tip of young elongating root hairs using a laser-mediated procedure allowing root hair selection (Wang *et al.*, 2019). Finally, the fact that *M. truncatula* root hairs can express two distinct types of K^+^-permeable outward conductance is further evidenced by the finding that one of these conductances can be also observed in protoplasts from root hairs of mutant plants that do not express *MtGORK*.

In summary, one of these two conductances is dependent on *MtGORK* functional expression and displays the same functional features as MtGORK when heterologously expressed in oocytes. It thus results from MtGORK activity. The second one, named fast-activating outward cationic conductance as in Wang *et al.* (2019), cannot be hypothesized to result from the activity of another Shaker channel. Indeed, within the *M. truncatula* Shaker family, *MtGORK* is the only outward channel gene. All the other genes belong to groups 1 to 4 (Supplemental Figure S1) and are thus likely to code for inwardly-or a weakly inwardly-rectifying channels that display activation upon membrane hyperpolarization. A simple hypothesis, based on the present knowledge, is that this fast-activating outward conductance corresponds to either a Cyclic-Nucleotide-Gated Channel (CNGC), a Glutamate Receptor (GLR) or an Annexin. The electrophysiological properties of the members from these families are still poorly characterized (Hedrich, 2012).

This K^+^-permeable fast-activating outward conductance is reminiscent of conductances reported in other cell types and plant species, for instance the NORC conductance characterized in barley root xylem parenchyma (Wegner and Raschke, 1994; Wegner and De Boer, 1997) and the weakly voltage-dependent non-selective cation conductance described in wheat roots (Davenport and Tester, 2000). However, this type of conductance was not reported in previous electrophysiological analyses carried out in root hairs from Arabidopsis (Ache *et al.*, 2000, Ivashikina *et al.*, 2001) and alfalfa (Bouteau *et al.*, 1999; Kurkdjian *et al.*, 2000). This might indicate that the levels of expression or activity of this conductance in root hairs is strongly dependent on the plant material, growth conditions and plant species, or is specific of *M. truncatula* when compared with Arabidopsis and alfalfa.

In most wild-type root hair protoplasts (*ca*. 80% of the protoplasts), the membrane outward conductance to K^+^ was strongly dominated by either the MtGORK or the fast-activating conductance, with rather similar frequencies, and current features revealing that the two conductances were simultaneously active could be detected in only ca. 20% of the protoplasts. Hence, it is likely that heterogeneities in the electrical properties of the root hair plasma membrane exist, at least transiently, amongst young root hairs. Such heterogeneities might be related to those of root hair tip responses to Nod Factors (NF), since NF treatment do not result in an alteration of tip growth in the same way in every young root hair (Esseling *et al.*, 2004).

### Comparison of the roles of MtGORK and AtGORK in control of stomatal aperture

In guard cells, the membrane outward K^+^-permeable conductance was dominated by MtGORK (Figure 2). It is interesting to note that no fast-activating outward conductance similar to that expressed by root hairs was detected in guard cells (Figure 2B), and that the absence of MtGORK activity in the mutant plants did not appear to be compensated by a conductance of this type (Figure 2B and 2C).

In Arabidopsis guard cells, AtGORK dominates the membrane outward conductance to K^+^ and mediates the depolarization-induced potassium release involved in stomatal closure (Ache *et al.*, 2001). Mutant plants harboring a KO mutation in *AtGORK* displayed slower closure kinetics, when compared with wild-type plants, resulting in impaired control of transpirational water loss (Hosy *et al.*, 2003). The data shown by Figure 3 reveal that the role of AtGORK in Arabidopsis guard cell upon stomatal closure is played by MtGORK in *M. truncatula*. Furthermore, they suggest that the control of leaf transpirational water loss and the contribution of the Shaker outward conductance to this function are stronger in *M. truncatula* than in Arabidopsis (dotted lines in Figure 3).

### Comparison of the roles of MtGORK and AtSKOR in K^+^ translocation to shoots

The *mtgork* KO mutation was found to be without any significant effect on the shoot K^+^ content in non-inoculated plants (Figure 4A), and to result in a slight reduction of this content in plants inoculated with the symbiotic partner *S. meliloti* (Figure 4B). Also, the concentration of K^+^ in the xylem sap driven by root pressure (after shoot excision) and the volume of exuded sap were not significantly affected by the *mtgork* mutation (Figure 5). Thus, absence of outward Shaker channel activity in *M. truncatula* root stele poorly affects K^+^ translocation towards the shoots, especially when compared with the corresponding results reported in Arabidopsis, where the absence of outward Shaker channel activity due to a KO mutation in *AtSKOR* results in a reduction in shoot K^+^ content and K^+^ concentration in the xylem sap by about 50% (Gaymard *et al.*, 1998). This indicates that MtGORK poorly contributes to K^+^ translocation towards the shoots, in contrast to AtSKOR, or that its absence in the mutant plants is efficiently compensated by other types of K^+^-permeable conductances. Non-Shaker K^+^-permeable outward conductances have been identified in xylem parenchyma cells from barley (Wegner and Raschke, 1994; Wegner and De Boer, 1997).

The K^+^ concentration of the collected xylem sap samples was about 5 times higher in the inoculated than non-inoculated plants (Figure 6). However, the shoot K^+^ contents were larger only by about 1.5 times in the former than in the latter plants (Figure 5). Furthermore, in the same experimental conditions, the shoot biomass was lower by about 2 times in the former than in the latter plants (Supplemented Figure S2A and S2B). This suggests that the flux of recirculated K^+^ ions from shoots to roots *via* the phloem sap is larger in symbiotic conditions. A larger flux of phloem sap towards the roots would provide sugars to functioning nodules. MtGORK, which is expressed in the nodule vasculature (Figure 4), may thus contribute to the recirculation towards the shoots of K^+^ ions arriving in nodules via the phloem sap. This hypothesis is consistent with the fact that, when inoculated, WT plants displayed (slightly) higher shoot K^+^ contents than mutant plants (Figure 5).

### Role of MtGORK in root hairs and early transduction of Nod Factor signal

In Arabidopsis root hairs, *AtGORK* encodes the typical K^+^-sensitive voltage-gated outwardly-rectifying conductance (Ache *et al.*, 2000), without any significant contribution of *AtSKOR* (Ivashikina *et al.*, 2001). AtGORK has been hypothesized in this cell type to be involved in control of cell turgor and membrane potential. In connection with this latter function, it has also been hypothesized to play a role in electrical signals (transient depolarization and changes in K^+^ fluxes) such as those induced by elicitor treatments (Ivashikina *et al.*, 2001). It should however be noted that none of these hypotheses has received direct support from reverse genetics approaches so far, and thus that the role of AtGORK in Arabidopsis root hairs is still unclear.

Here we show that MtGORK contributes to the repolarization of the root hair cell membrane following the NF induced depolarization. The repolarization still occurred in absence of MtGORK activity, but at a slower rate, by about two times. Thus, these results indicate that MtGORK plays an important role in the electrical signal triggered by NF perception by contributing to the membrane repolarization. However it is not the only electrogenic transport system involved in the repolarization since this process remains, although at a slower rate, in mutant root hair devoid of MtGORK conductance. The simplest hypothesis is that the fast-activating outward conductance identified in root hairs besides MtGORK (Figure 7) plays a role in the repolarization and can compensate for the absence of MtGORK activity in mutant plants.

We checked whether the slower repolarization of the root hair cell membrane in the mutant plants affected subsequent steps of plant engagement in the symbiotic interaction with *S. meliloti*. Comparison of the number of infection threads in WT and mutant plants grown *in vitro* on agar plates in presence of *S. meliloti* did not reveal any significant consequence of the *mtgork* mutation. Nodule production was also similar in the two types of plants when grown *in vitro*. Thus, these results suggest that the kinetics of membrane repolarization is not a crucial component of the signaling pathway leading to the symbiotic interaction of *M. truncatula* with *S. meliloti*.

## Conclusion

The most striking difference between MtGORK, representative of all the plant outwardly-rectifying Shaker channels characterized so far, and the fast-activating K^+^-permeable outward conductance, appears to be the rectification capacity. The fast-activating conductance is weakly rectifying and thus can allow K^+^ influx when the electrochemical gradient of this cation across the plasma membrane is inwardly directed, while the strong regulation of MtGORK by both the voltage and the external concentration of K^+^ ensures that the permeation pathway remains closed when the K^+^ electrochemical gradient is inwardly directed, so that these channels are strictly dedicated to K^+^ secretion. Based on the present knowledge, all plant species whose genome has been sequenced possess at least one Shaker channel of this type (Véry *et al.*, 2014). While some species possess 4 genes encoding such channels, like grapevine or poplar (Véry *et al.*, 2014), a single one is sufficient in other species. It is interesting to note that, in species displaying a single outward Shaker channel like *M. truncatula*, this channel can display a rather broad expression pattern, suggesting that it might be involved in the various functions involving its different orthologs in species that harbor several channels of this type. Within the framework of this hypothesis, the fact that the absence of MtGORK channel activity poorly affects K^+^ translocation towards the shoots, when compared with the effects of the corresponding mutation in Arabidopsis, would result from compensation in some tissues of the absence of MtGORK by other types of conductances in the mutant plants. It should be noted that, in root hairs in contrast to guard cells, the fast-activating K^+^-permeable conductance appears to be able to compensate the absence of MtGORK conductance in mutant plants. Despite such possibilities of redundancy and compensation, the fact that all plant species possess at least one outward Shaker channel gene indicates that K^+^ channels displaying a strict outward rectification provide important services in some environmental conditions.

## MATERIAL AND METHODS

### Plant material and plant growth

*M. truncatula* (ecotype Jemalong A17) seeds were scarified with sulfuric acid (99%) for 10 min, rinsed and sterilized in 6% sodium hypochlorite solution for 3 min. After 3 h imbibition in sterile water, seed coats were removed and seeds were transferred onto 1% agar plates in Petri dishes, which were turned upside down (agar up) and remained in the dark at 4°C for at least 48 hours in order to break dormancy and obtain synchronization of germination. The plates were then transferred at 21 C for 16 to 24 h for germination. The radicles were then about 2 cm long.

For *in vitro* culture, germinated seedlings were transferred onto a sterile sheet (12 × 8.5 cm) of chromatography paper (Rogo-Sampaic, France) laid on solid Fahraeus agar medium (modified from Vincent, 1970) in a Petri dish (12 × 12 cm, for 10 seedlings). The medium contained 10 g L^−1^ of purified agar (Euromedex, https://web.euromedex.com/) and 0.5 mM MgSO_4_, 0.7 mM KH_2_PO_4_, 0.8 mM Na_2_HPO_4_, 1 mM CaCl_2_, 20 µM Fe-citrate and 0.1 mg.L^−1^ of MnSO_4_, CuSO_4_, ZnSO_4_, H_3_BO_3_ and Na_2_MoO_4_, pH 7.5 (adjusted with KOH). The lower part of the plate was wrapped in aluminum foil to avoid detrimental effects of light on roots. The plate was placed in a quasi-vertical position in a growth chamber (70% relative humidity, 70 μE.m^−2^.s^−1^ light intensity) with a photoperiod of 16 h light (25 °C) and 8 h dark (21 °C) for 5 days.

Germinated seedlings grown on the agar plates for 2 further days were transferred on sand-vermiculite mixture (3:1, v:v; ca. 1 L per plant in 10 L containers) or sand-compost (3:1, v:v; ca. 1.5 L per plant in individual pots) and grown in a growth chamber (70% relative humidity, 16 h light, 300 µE.m^−2^.s^−1^ light intensity, 25°C, and 8 h night, 21°C) or in greenhouse, respectively. They were watered twice a week, alternatively with water or Fahraeus solution).

Composite plants were generated according to the protocol of Boisson-Dernier *et al.* (2001) using the electrocompetent *Agrobacterium rhizogenes* strain ARqua1 harboring the transcriptional *GUS* construct (2 kb *gork* promoter sequence cloned in the *pGWB3* vector from Gateway system) (Nakagawa *et al.*, 2007).

### Mutant and control wild-type plants

*M. truncatula* mutant line NF9352 was identified (BLASTn of the *GORK* genomic sequence in the Noble website) in the Noble collection (http://medicago-mutant.noble.org/mutant/) as harboring a *Tnt-1* retrotransposon insertion in the *MtGORK* gene (Supplemental Figure S1D). We named the mutation resulting from this insertion *mtgork*. A F1 plant hemizygous for the *mtgork* mutation was amplified. PCR genotyping experiments on the F2 progeny identified plants either homozygous for the *mtgork* mutation or displaying a wild type genotype at this locus and thereafter named control WT plants. Both types of plants were amplified for F3 progeny.

### Rhizobial strain and plant inoculation

The rhizobial strains used for *M. truncatula* inoculation were *Sinorhizobium meliloti* Rm1021, Rm1021 DsRed and Rm2011 LacZ. Bacteria were grown in 5 g.L^−1^ Bacto tryptone, 3 g.L^−1^ yeast extract, 6 mM CaCl_2_, pH 7.2 (TY/Ca medium), supplemented with the appropriate antibiotic: 50 µg.mL^−1^ streptomycin for Rm1021 and Rm1021 DsRed and 10 µg.mL^−1^ tetracycline, centrifuged, resuspended and washed in Fahraeus medium. Aliquots of the final bacterial suspension (OD_600_ ~ 10^−2^) were directly laid over apices of the germinated seedlings. Sand-vermiculite and sand-compost mixtures were also inoculated (*ca.* 10 mL of the rhizobial suspension for 1 L of soil).

### Plant K^+^ contents and K^+^ translocation to shoots via the xylem sap

Plants were grown on sand-vermiculite mixture for 8 weeks in growth chamber. Shoots and roots were collected, dried and weighted (DW). Ions were extracted with 0.1 N HCl and assayed (flame spectrophotometry). The shoots of plants grown in parallel, in the same conditions and for the same time, were excised below the first leaf, at about 1 cm above the soil. For each plant, about 30 min after shoot excision, the root extremity was introduced into a plastic tube (Eppendorf type) through a hole pierced at the tube bottom and sealed with silicon paste to collect exuded sap for 24 h. The volume and K^+^ and Ca^2+^ concentrations of the exuded sap were measured.

### Two-electrode voltage clamp characterization of MtGORK in *Xenopus* oocytes

The coding sequence of *MtGORK* was amplified by PCR, cloned into the pGEM-Xho vector (derived from pGEMDG; D. Becker, Würzburg) downstream from the T7 promoter and between the 5′- and 3′-untranslated regions of the Xenopus β-globin gene. Capped and polyadenylated copy RNA (cRNA) were synthesized *in vitro* (mMESSAGE mMACHINE T7 kit, Ambion). Oocytes, isolated and handled as described previously (Véry *et al.*, 1995), were injected with *ca.* 30 ng of *MtGORK* cRNA (ca. 1 ng.nL^−1^) or with 30 nl of diethyl-pyrocarbonate-(DEPC) treated water for control ("water injected") oocytes. They were then kept at 18 °C in ND96 solution (96 mM NaCl, 2 mM KCl, 1.8 mM CaCl_2_, 1 mM MgCl_2_, 2.5 mM sodium pyruvate, and 5 mM HEPES-NaOH, pH 7,4) supplemented with 0.5 mg L^−1^ of gentamycin until voltage-clamp recordings. Whole-oocyte currents were recorded using a two-electrode voltage-clamp technique 1–2 d after cRNA injection. All electrodes were filled with 3 M KCl. The external solution bathing the oocyte was continuously percolated during the voltage-clamp experiment. All bath solutions contained a background of 1 mM CaCl_2_ and 2 mM MgCl_2_, buffered with either 10 mM Hepes-NaOH or 10 mM Mes-NaOH, pH 7.5 or 5.6, respectively. This background solution was supplemented with either 100, 30, 10 or 3 mM KCl, the osmolarity of the solution being maintained constant by addition of N-Methyl-D-glucamine (NMDG), or with 100 mM CsCl, RbCl, NaCl or LiCl. To extract MtGORK-mediated currents from total oocyte currents, mean currents recorded in water-injected control oocytes from the same batch in the same ionic conditions were subtracted from those recorded in MtGORK-expressing oocytes.

### Leaf transpirational water loss

Plants of WT and *mtgork* lines were grown in greenhouse in compost for 6 weeks. Then, leaves were excised and exposed to desiccation by placing them on trays on the lab bench at room temperature. Leaves were weighed (FW: fresh weight) at different time points. Transpirational water loss was determined from the difference between the weight at each time point and the initial weight.

### Protoplast isolation and patch-clamp analyses

Root hair protoplasts were obtained by enzymatic digestion as previously described (Wang *et al*., 2019) and stored in ice until patch-clamp measurements. For guard cell protoplast preparation, epidermal strips were peeled off from the abaxial surface of 6 to 7 leaves using forceps, and cut into small pieces. The enzymatic treatment was performed for 1h and 40 min at 28°C in a solution containing 1% (w/v) cellulase RS, 0.1% (w/v) pectolyase Y23, 1% (w/v) BSA, 1 mM CaCl_2_, 2 mM ascorbic acid, 1 mM Mes, and 450 mM D-mannitol, its pH being adjusted to 5.7 with KOH. Then, the released protoplasts were collected by filtering through a 40 µm nylon mesh and washed with the conservation medium twice and stored in ice until patch-clamp measurements. The conservation medium contained 1 mM CaCl_2_, 2 mM ascorbic acid, 1 mM Mes, and 500 mM D-mannitol, its pH being adjusted to 5.7 with KOH.

Patch-clamp experiments were performed in the whole-cell configuration. Patch-clamp pipettes were pulled using a DMZ-Universal Puller (Zeitz-Instruments GmbH, Germany) from borosilicate capillaries (GC150F-7.5; Phymep, France) and fire polished (by the DMZ-Universal Puller). Microelectrode resistance was about 10 and 14 MOhms for patch clamping root hair and, respectively, guard cell protoplasts. A reference Ag/AgCl half-cell completed the circuit. The patch clamp amplifier was an Axopatch 200B (Axon Instruments Inc., USA). Whole-cell currents were measured at least 5 minutes after seal formation. Data were sampled at 1 kHz. The Clampex module of the pClamp9 software (Axon Instruments Inc., USA) was used for data acquisition. Analysis was performed using the Clampfit module of pClamp10 and SigmaPlot 11 (Systat Software Inc., USA). Liquid junction potentials were corrected.

### Membrane potential measurements

*M. truncatula* seedlings were grown on agar medium (10 g L^−1^ of purified agar in a solution, named bath background solution, bbs, containing 0.1 mM KCl, 0.1 mM CaCl_2_, 0.1 mM NaCl and 2 mM MES-Bis-Tris-Propane, pH 6.5) for 2 days. Roots were excised and fixed in a plexiglass chamber filled with bbs. The chamber was percolated with bbs for 15 min (recovery treatment) before root impalement. Impalement micro-electrodes, with a tip diameter of approximately 0.5 µm, were pulled from borosilicate glass capillaries (GC200F-10, Harvard Apparatus, http://www.harvardapparatus.com) and back-filled with 3 M KCl. Microelectrodes were connected via an Ag/AgCl pellet to an HS-2 ⋅ 0.1L probe of an Axoprobe 1A electrometer (Axon Instruments). The reference comprised a combined glass pH electrode (filled with 3 M KCl) placed in the chamber downstream of the root. The micro-electrode was placed at the root surface, at about 0.5 cm from the tip, in front of young developing root hairs, using a manually operated micro-manipulator (Narishige, http://narishige-group.com). Subsequently, the vertical position of the root chamber was adjusted using a micro-elevator (IT6D CA1, Microcontrole, http://www.newport.com), allowing precise penetration of the micro-electrode into an epidermal cell at 10-40 µm below the root surface. During impalement, the bath solution was continuously refreshed. The steady-state membrane potential was successively measured in 5 external solutions: bbs, bbs + 10^−6^ M Nod factor, bbs, bb + 30 mM KCl and bbs again. The whole protocol was achieved within less than 30 min. The recording was discarded when the membrane potential values got in the 3 bbs successively perfused during the protocol were not consistent together.

### Promoter fusion and histochemical localization of GUS Activity

A 2-kb DNA fragment upstream of the starting ATG of *MtGORK* gene (*Medtr5g077770*), was amplified by PCR using gene-specific primers (pSKOR2kb-F1: CACTCCTTAGCAAATAGCAAAAATTA and pSKOR2kb-R1: GAAATTAATTAATTAACCTATCCTTAGAAG). Composite *M. truncatula* plants were obtained by transformation with *Agrobacterium rhizogenes* ArquaI strain as previously described (Andrio *et al.*, 2013). Healthy composite plants were transferred onto new plates containing modified Fahraeus medium without nitrogen and kanamycin. Plants were inoculated with *S. meliloti* 3 d after transfer.

Transgenic roots were stained with GUS assay buffer as previously described (Andrio *et al.*, 2013). Roots from at least 20 plants from three biological experiments were examined. Roots and nodules were fixed in 1% glutaraldehyde and 2% formaldehyde in 0.05 M phosphate buffer (pH 7), washed, dehydrated, and embedded in Technovit 7100 according to the manufacturer’s instructions. Fifty-micrometer-thick vibroslices were obtained with a HM560V Vibratome (Leica RM 2165) and visualized with an Olympus BH-2 microscope using dark-field optics.

## ACKNOWLEDGEMENTS

This work was supported in part by an ANR grant (ANR-11-BSV7-010-02) (to HS and AAV), a doctoral grant of the Government of Nouvelle Calédonie (to AD) and the French Ministry of Research (to JT).

## SUPPORTING INFORMATION

Additional Supporting Information may be found in the online version of this article.

**Supplemental Figure S1. MtGORK of the Shaker channel family of *Medicago truncatula*.**

(A) Phylogenetic relationships between Shaker polypeptides from *Arabidopsis thaliana* and *Medicago truncatula*. The plant Shaker family comprises 5 groups (see main text). Shaker sequences from Arabidopsis were obtained from the TAIR website (http://www.arabidopsis.org/). A homology search was carried out against the *M. truncatula* protein sequence bank (MT4.0v2) using the BLAST (Basic Local Alignment Search Tool) program, the BLOSUM62 matrix (BLOcks SUbstitution Matrix) and a threshold E (or E-value) equal to 10^−3^. The unrooted phylogenetic tree was generated with PhyML software (http://www.atgc-montpellier.fr/phyml/binaries.php) using the maximum-likelihood method and 1000 bootstrap replicates in Seaview application (http://doua.prabi.fr/software/seaview). Arabidopsis Shaker polypeptide sequences were first aligned with Muscle (http://www.drive5.com/muscle/), then treated with Gblocks in Seaview program for alignment curation. The phylogenetic tree was drawn with Dendroscope (http://ab.inf.unituebingen.de/software/dendroscope/). Bootstrap values (as percentages) are indicated at the corresponding nodes. The scale bar corresponds to a distance of 10 changes per 100 amino acid positions.

(B) Structure of plant Shaker channels. The channel hydrophobic core comprises 6 transmembrane segments, named S1 to S6. S4 (the so-called voltage sensor) contains positively charged residues and confers sensitivity to the electric field in the membrane (and thus to the transmembrane voltage). P: pore domain. Shaker functional channels are tertrameric proteins, the 4 P domains being assembled in the center of the tetrameric structure where they structure the K^+^ permeation pathway (pore). Four large domains can be identified in the cytosolic region downstream S6: a C-linker domain, a cyclic nucleotide binding domain, an ankyrin domain (not present in every plant Shaker channel but present in MtSKOR, AtSKOR and AtGORK), and a K_HA_ domain. Role of these domains in plant Shaker channels: see Daram *et al.*, 1997; Nieves-Cordones *et al.*, 2014).

(C) Sequence alignment of MtGORK and the Arabidopsis AtSKOR Shaker in the P domain and S6. Asterisks denote residues shown to be involved in the channel sensitivity to the external concentration of K^+^ in AtSKOR (Johansson *et al.*, 2006).

(D) Schematic diagram of the *MtGORK* gene structure indicating the site of insertion of the disrupting TNT1 retrotransposon in the Jemalong A17 *M. truncatula* line named NF9352. Same abbreviations as in panel B. Boxes: exons.

**Supplemental Figure S2. Mutant plants homozygous for the *mtgork* mutation display similar shoot and root biomass as wild type plants.**

Plants homozygous for the *mtgork* mutation (mutant) or displaying a wild type genotype for this mutation (WT) were compared with respect to biomass production in different conditions. (A and B) Non-inoculated (A) or inoculated (B) plants grown for 8 weeks on vermiculite-sand mixture. Plants were transferred onto sand-vermiculite mixture 3 days after germination. When inoculated (B), inoculation (with *S. meliloti* strain 1021) was achieved after 7 days of growth. Shoot and root were collected for biomass measurements (dry weight: DW) after 8 weeks of growth in growth chamber. Means ± SE; n = 12.

(C) Inoculated plants grown *in vitro* for 3 weeks. Germinated seedlings were transferred onto Fahraeus agar medium in Petri dishes and inoculated with *S. meliloti* strain 1021 DsRED. Plant dry weight was measured 21 days post-inoculation (means ± SE; n = 24).

(D) Inoculated plants grown for 2, 3 or 4 weeks on compost. Germinated seedlings were transferred onto compost on growth chamber and inoculated one week later with *S. meliloti* strain 1021. Plants were collected for dry weight measurements at 7, 14 and 21 dpi.

