## Supplementary figures and images for "*Medicago truncatula* possesses a single Shaker outward K^+^ channel: functional characterization and roles *in planta*"

### Supplemental Figure 1

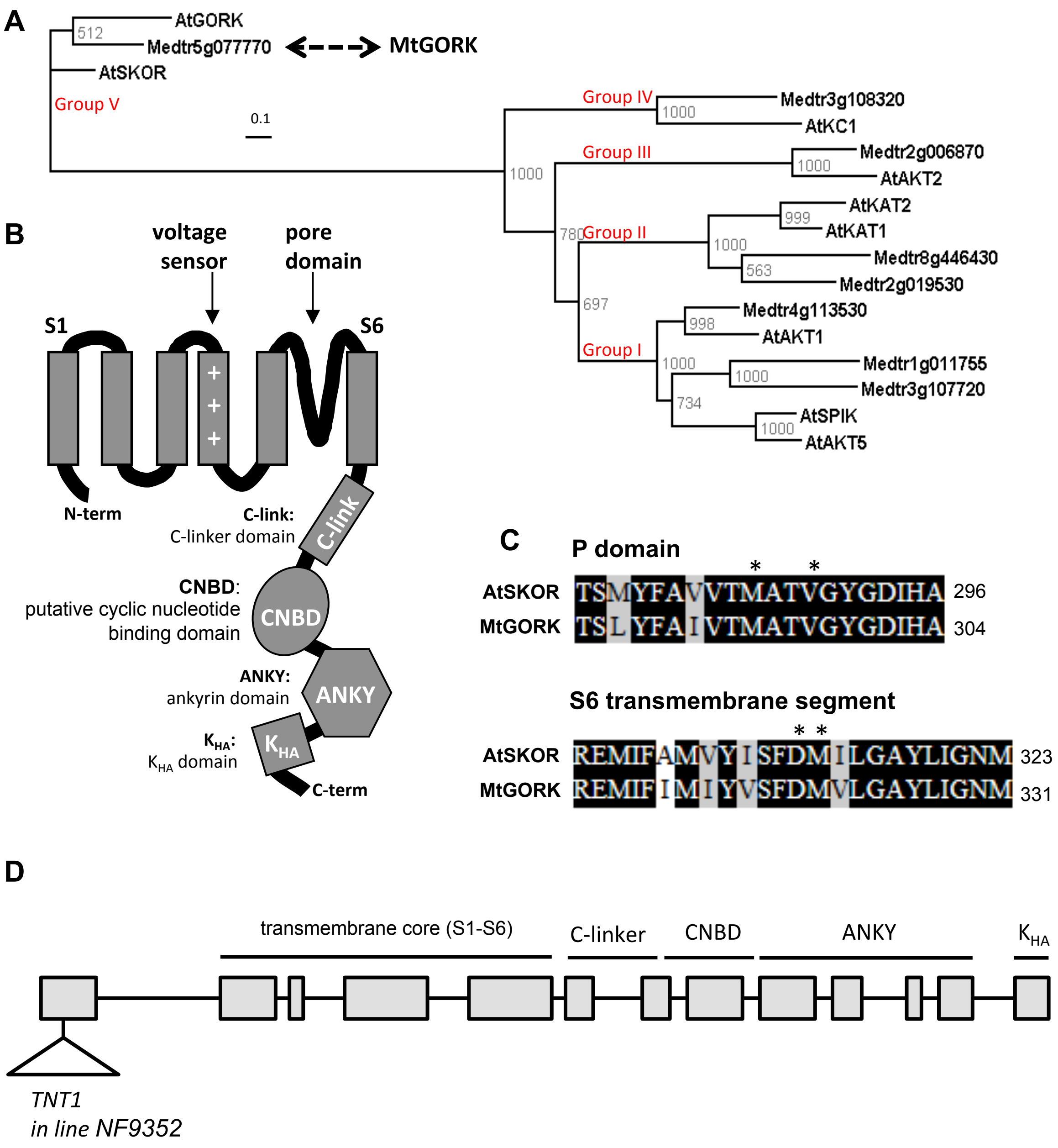


**Supplemental Figure 1**

### Supplemental Figure 2

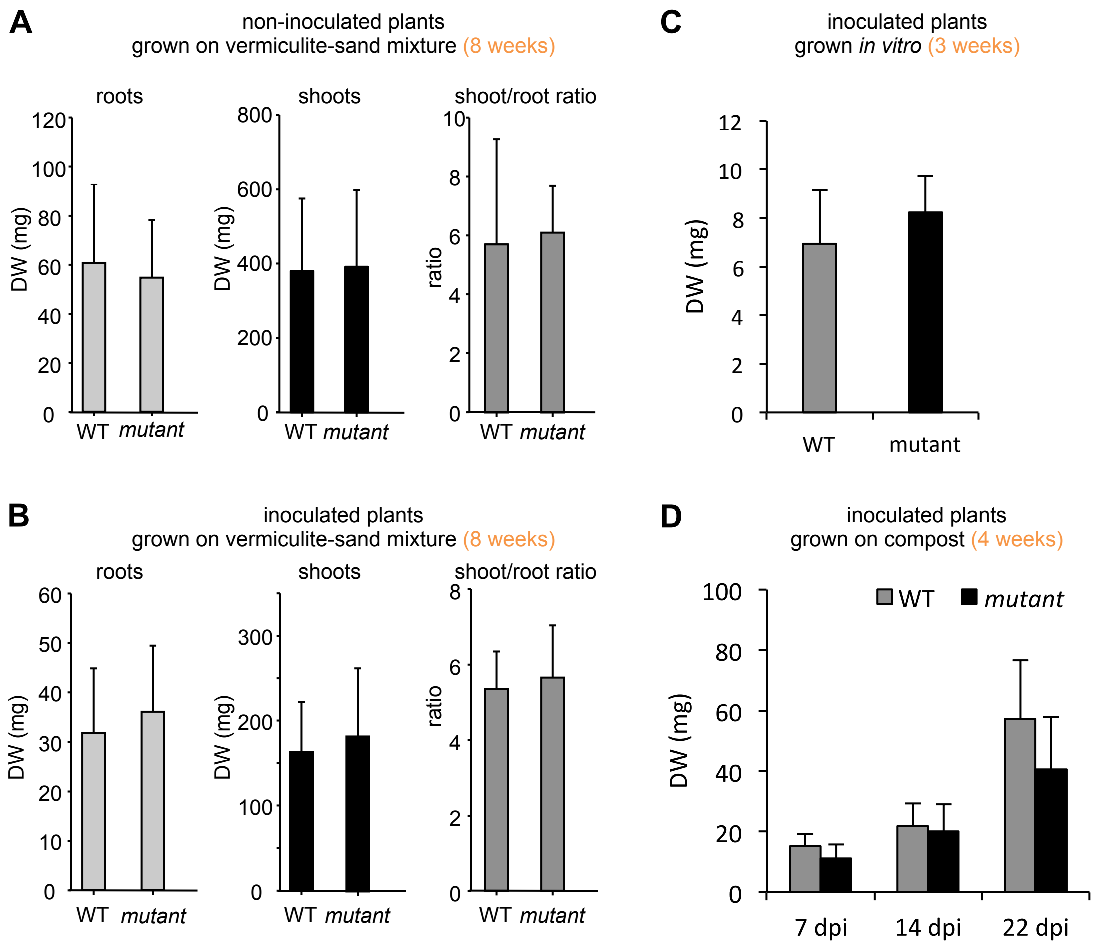


**Supplemental Figure 2**
